# Contribution of genome scale metabolic modeling to niche theory

**DOI:** 10.1101/2021.07.21.453190

**Authors:** Antoine Régimbeau, Marko Budinich, Abdelhalim Larhlimi, Juan Jose Pierella Karlusich, Olivier Aumont, Laurent Memery, Chris Bowler, Damien Eveillard

## Abstract

Standard niche modeling is based on probabilistic inference from organismal occurrence data but does not benefit yet from genome-scale descriptions of these organisms. This study overcomes this shortcoming by proposing a new conceptual niche that encompasses the whole metabolic capabilities of an organism. The so-called metabolic niche resumes well-known traits such as nutrient needs and their dependencies for survival. Despite the computational challenge, its implementation allows the detection of traits and the formal comparison of niches of different organisms, emphasizing that the presence-absence of functional genes is not enough to approximate the phenotype. Further statistical exploration of an organism’s niche sheds light on genes essential for the metabolic niche and their role in understanding various biological experiments, such as transcriptomics, paving the way for incorporating better the genome-scale description in ecological studies.

---

A hundred years ago, seminal studies introduced the general idea of a niche that still motivates current ecological investigations. In 1917, Joseph Grinnell proposed one of its first definitions by declaring the niche as the environmental conditions needed by a given species to survive (1). However, because such a description did not consider the impact of the species on its environment, Charles S. Elton (2) proposed a complementary description examining the niche as the place of the species in its biotic environment. We have to wait until 1957 for G. E. Hutchinson to publish his *Concluding Remarks* (3) that explored a new formalism that could embed both definitions. He proposes a niche space as an n-dimensional space, where each axis describes an environmental variable. The set of conditions allowing a species to survive defines its niche, forming an n-dimensional volume in the niche space. This definition is referred to as the *fundamental niche*. It aims to reason on the biological system requirements and to highlight its impact on its environment. Nevertheless, modeling such n-dimensional volume is challenging because of the nature of biological data. To overcome these limitations, many heuristics, leading to different definitions of the niche, are proposed. However, these numerous formalizations contribute to the complexification of the niche concept (4).

In parallel, high-throughput technologies have changed the global perception of a biological system and fostered the use of DNA sequencing techniques and molecular abstractions in microbial ecology (5). Meanwhile at the interface of computer science, mathematics, and molecular biology the new field of systems biology emerged (6). This discipline’s primary goal is to extract emerging properties from biological systems depicted by high-throughput molecular experiments. Increasing computing capacity and large dataset availability enabled systems biology to extend its modeling application domain from small size reductionist networks to ecosystems (7–10). Nowadays, biological systems are analyzed by their gene content, allowing metabolic network reconstruction that attempts to predict the metabolic phenotype (i.e., biochemical capabilities of an organism) from the genotype (11). These metabolic predictions from omics data have shown significant successes in biotechnology (12–15). However, these predictions assume the biological system to adopt optimal behaviors such as growing at their maximal rate, which is not suited for the niche concept, where organisms show their plasticity to survive, not their ability to overgrow.

Here we proposed a novel computational framework where we extract the niche of an organism based on its metabolic network. We first use the quantitative description of this *metabolic niche* to recover biological features of *Escherichia coli* such as conditions for aerobic or anaerobic growth

. We then extend our computations to numerous prokaryotes, exhibiting metabolic niche inclusion and its putative link with ecotype. Finally, we investigate the metabolic niche of *Phaeodactylum tricornutum* showing the importance of particular reactions and pathways in the survival of the organism. Notably, we show that we can gather different kinds of omics data around our theoretical framework for improving our understanding of previous biological results in light of the ecological success of diatoms.

## Results

### Rationale of the metabolic niche and its application on a state-of-the-art metabolic model

The metabolic niche is a new computational result estimated from the sole genomic composition of an organism. Contrary to other niche estimations (16), our modeling does not consider statistical distributions over a range of parameters but rather a theoretical phenotype. From genomic data (Fig. 1a), metabolic network reconstruction techniques infer an organism’s metabolic abilities. Reactions intertwine with products of reactions that are substrates of others, forming a metabolic network (Fig. 1b). Some metabolites are constituents of the biomass reaction. This synthetic reaction models the growth rate of the organism. Exchange reactions are responsible for the organism’s inter-action with its environment, defining the limits of the biological system. The set of all biological constraints, such as (i) the inter-connectivity of reactions, (ii) the stoichiometry, and (iii) a classical steady-state approximation, describes a solution space (Fig. 1c). From this solution space, state-of-the-art systems biology approaches focus on distinct points of interest, generally minimizing or maximizing one particular flux (e.g., the red dot in Fig. 1c maximizing the flux through the biomass reaction). The fundamental niche focuses on the species’ interaction with its environment and survival (Fig. 1d). Projected in a metabolic modeling framework, considering the niche space implies abstracting the network’s inner reactions and focusing on the exchange reactions while maintaining a minimal growth rate (Fig. 1e). The former metabolic solution space is therefore further constrained by a death rate. Hence, the corresponding space describes all fluxes combinations that satisfy the fundamental niche constraints, so-called the metabolic niche (Fig. 1f).

**Fig. 1.**
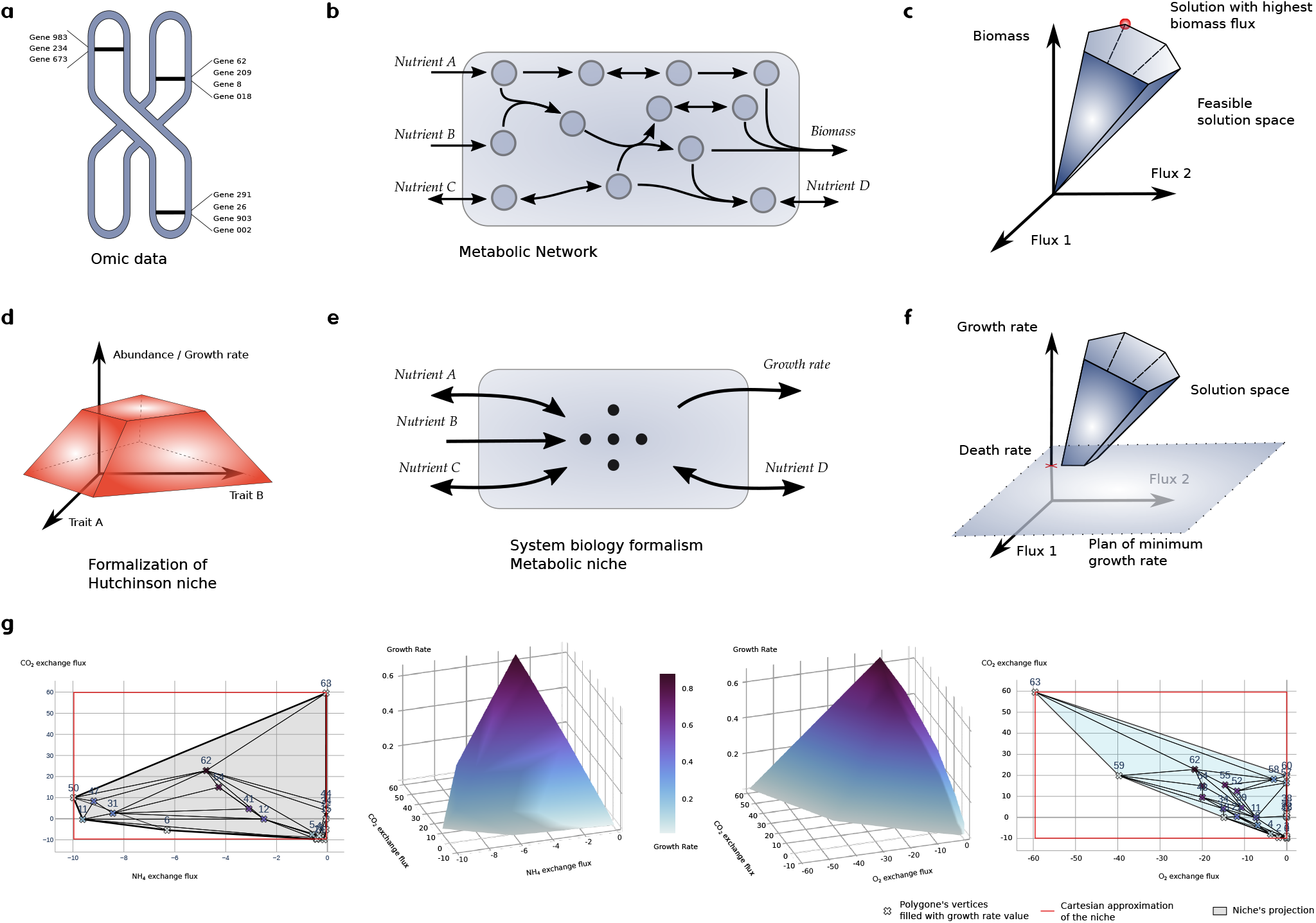
Formalizing the metabolic niche from omic knowledge and application on E. coli. From the genomic content of an organism (**a**), we could infer the corresponding encoded catalytic proteins. These proteins support metabolic reactions that interplay (i.e., products are substrates for others). The resulting metabolic network (**b**) integrates all the physiological abilities of one species. Projected into a constraint-based paradigm, solving the corresponding linear problem depicts a solution space (**c**) that describes all the fluxes in each metabolic reaction that satisfy all given constraints. Ecological traits define the axes of Hutchinson’s niche space. The niche volume (**d**) guarantees that the species survives as long as its environmental conditions belong to this volume. If we apply this formalism to the metabolic network, we abstract the organism’s inner mechanism and focus only on its exchange reactions (**e**). Using constraint-based modeling on this new system, we have a new constraint, which is the survival of the species; that is, the flux through the growth rate reaction needs to be at least as high as the death rate of the species (**f**). Therefore, a formal description of the metabolic niche explores the solution space in which axes (or traits) are exchange reaction fluxes. When applied on *E. coli*, we reduced the number of axes of the niche space to allow visualization of the volume of its niche (**g**)

For the sake of application, a metabolic niche was estimated for the core *Escherichia coli* metabolic network. An estimation of the maximum growth rate via Flux Balance Analysis (17) is 0.874 mmol.gDW^−1^.*h*^−1^. The metabolic niche was estimated for a death rate equals to 0.01 mmol.gDW^−1^.*h*^−1^ and on six dimensions, or traits, that are exchange reactions for the following metabolites: CO_2_, O_2_, H_2_O, NH_4_, phosphate, and glucose. The death rate value is herein strictly arbitrary, and results may vary for other values. For instance, a value of 0.7 would shed light on high growth rate behavior, whereas a lower value would add behaviors representing less than 15% of the flux range through the biomass. A finer parameterization of this rate is necessary to represent biotic interactions better, such as predation, but does not interfere in the fundamental niche definition.

The topology of a metabolic niche is complex. We could describe its volume with more than 210 vertices. In favor of representation, we clustered close vertices, dividing by three their number (64 vertices) while maintaining more than 95% of the niche volume for the represented axes. For the sake of simplicity, we represent the metabolic niche (a blue-grey area) and its vertices in the CO_2_ vs. ammonium and CO_2_ vs. O_2_ exchange spaces (Fig. 1g). As a companion illustration, we also represented the metabolic niche in the front of the growth rate as a 3D shape to picture each environmental condition’s theoretical growth.

For approximating the state-of-the-art niche space in the context of the metabolic network, we performed a Flux Variability Analysis (FVA) (17) on the same exchange reactions with a minimum flux through the biomass reaction set to 0.01 mmol.gDW^−1^.*h*^−1^1 to mimic a death rate. It defined the feasible range of fluxes for each trait (Fig. 1g red lines). As a modeling validation, estimated ranges of fluxes embed the projection of the metabolic niche. The volume defined by the Cartesian product of each segment of the feasible range is called the Cartesian niche. The Cartesian niche volume is more than 86 × 10^6^ units in the six dimensions, whereas the metabolic niche volume is less than 72 × 10^3^. Thus, it occupies less than 1‰ of the approximate Cartesian niche, emphasizing all the constraints that are not taken into account in the Cartesian niche approximation, in particular those that define phenotypic traits. This support previous results on trait space occupied by plant compared to models (18).

The metabolic niche indicates an overall uptake of ammonia by *E. coli* (i.e., negative exchange fluxes), whereas CO_2_ could be produced (i.e., positive exchange fluxes) by respiration, or uptaken but in small amplitude for anaplerotic reactions and other carboxylation reactions. Flux distributions in this part of the metabolic niche are associated with lower uptake of oxygen (15 mmol.gDW^−1^.*h*^−1^), emphasizing the anaerobic growth conditions. As a biological validation, the metabolic niche description correctly shows lower maximal growth rates in anaerobic than in aerobic conditions conditions. Furthermore, the metabolic niche’s overall shape exhibits a positive relationship between O_2_ consumption and CO_2_ production that defines respiration ( i.e., a negative relationship between both fluxes), whereas the relationship between CO_2_ and ammonia is less obvious. This system’s characteristic is a straightforward consequence of considering whole intracellular biochemical reactions, leading to mechanistic interdependencies between uptake reactions, that by construction, Cartesian niches could not consider. Furthermore, the metabolic niche extracts essential numerical descriptors of *E. coli*, mainly physiological switches of regime in function of available nutrients. These switches between regimes are traits, usually identified after extensive and tedious experimental efforts.

### Comparing organisms via their metabolic niches

We arbitrarily defined traits for a systematic comparison of metabolic niches (i.e. exchange reactions for the following metabolites: NH_4_, SO_4_, H_2_S, glucose, NO_3_). We randomly selected 39 prokaryotic metabolic models (19) that share those traits. When possible, we associated each model with its habitat(20). We then compared the metabolic niche distance, a surrogate to the metabolic niche volume overlap, versus different state-of-the-art distances between organisms (Fig. 2a-c). Identical or included metabolic needs imply similar or included metabolic niches, and a metabolic niche-based distance equals to 0. Conversely, a distance equals to 1 emphasizes distinct niches and distinct metabolic needs. Each point describes a comparison between two bacteria. Colored points indicate bacteria from similar habitats (resp. blue, brown, and red for marine, soil, and host-associated). When removing inclusion cases, we show a strong positive correlation between Cartesian niche overlaps and metabolic niche overlaps (Fig. 2a; R: 0.82, p-value: 2.68 × 10^−181^, slope: 0.98). This result was expected as the Cartesian niche embed the metabolic niche. Furthermore, most of the comparisons are above the scatter plot’s diagonal. Several metabolic model comparisons are spread over vertical lines: showing the same Cartesian niche overlap but different metabolic niche overlaps, stressing an overall refinement in the distance brought by the use of the metabolic niche.

**Fig. 2.**
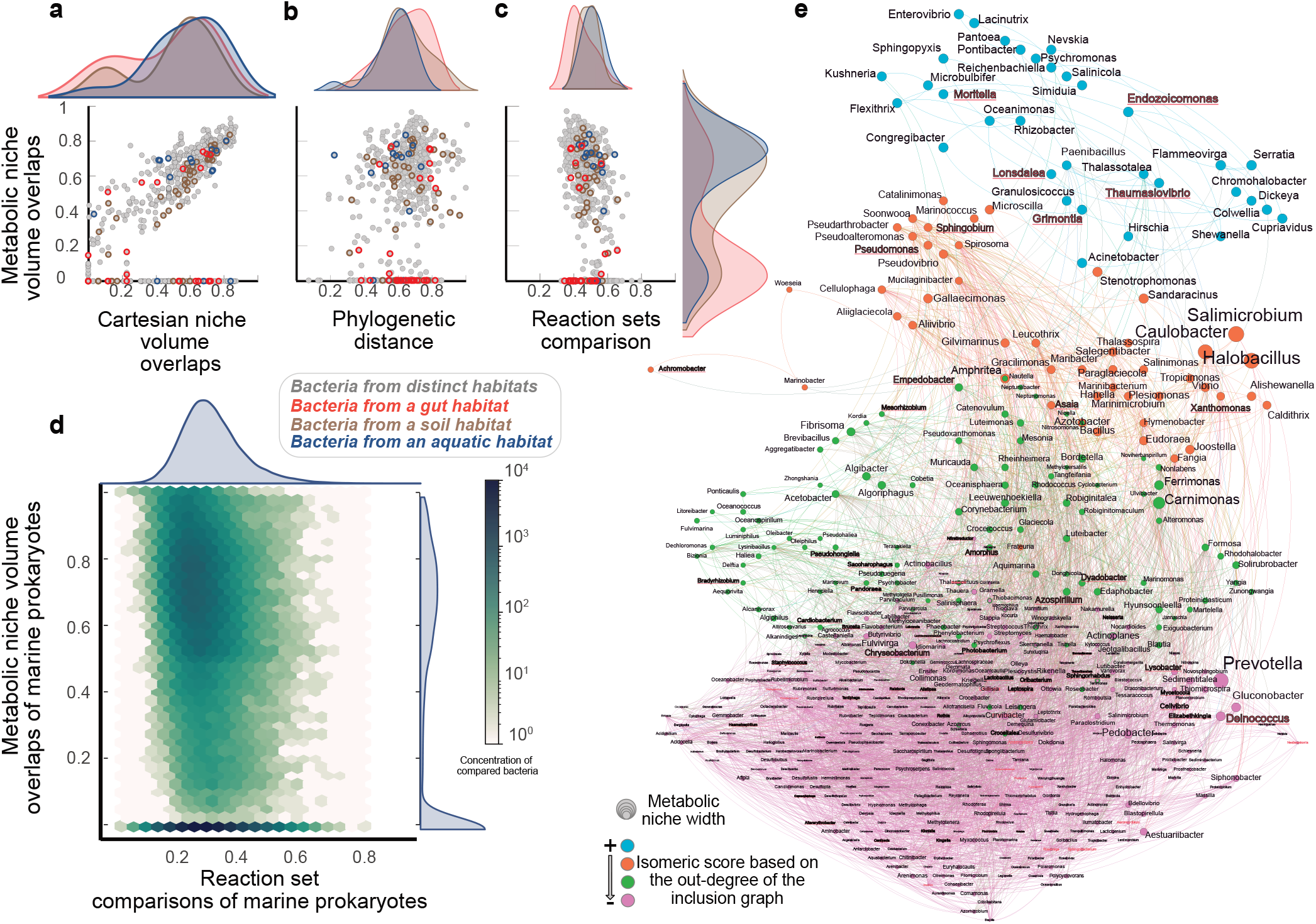
Comparison of the metabolic niche overlap with other pairwise distances and niche inclusion. **a**, comparison between metabolic niche overlaps and Cartesian niche overlaps estimated with a Jacquard distance. This comparison is performed for bacteria from different or common habitats. **b**, absence of a relationship between metabolic niche overlaps and pairwise phylogenetic distances based on 16S rRNA gene. **c**, absence of a relationship between metabolic niche overlaps and presence/absence of common reactions estimated via a Jaccard distance. **d**, absence of a relationship between metabolic niche overlaps and presence/absence of common reactions estimated for 502 marine bacteria metabolic models. Instead of a scatter plot, it shows concentrations for more than 10^5^ pairwise comparisons. For bacteria that show metabolic niche overlaps almost null, we investigate potential niche inclusion. **e**, metabolic niche inclusion graph of marine bacteria. The node size is proportional to metabolic niche volume, and an edge indicates a metabolic niche’s inclusion into another. The graph layout follows an isomeric distribution driven by the out-degree of the inclusion graph. Bacteria are divided into four categories based on the z-score from most embedded metabolic niche (i.e., turquoise) to the least (i.e., purple). Labels of bacteria known for being associated with a host are in red/bold and underlined.

Similarly, we compared the metabolic distance with the phylogenetic distance based on 16S rRNA sequence pairwise comparison (Fig. 2b). The comparison exhibits a decoupling between the metabolic niche and taxonomy as already shown for marine prokaryotes (21). Because seminal genomic studies extrapolate organismal functionalities from their genomic content (21, 22), we further compared the metabolic niche overlap with a genomic composition distance by computing pairwise organismal reactions sets’ Jacquard distances (Fig. 2c). In this context, high distance means low cardinality of the intersection of the sets. This straightforward metric stresses the topologies’ differences from a metabolic network perspective by emphasizing similar reactions between organisms but not their use. Our computations reveal no significant relationship between the metabolic niche overlap and the presence-absence of reactions, emphasizing that the exclusive metabolic abilities do not approximate the organismal metabolic needs or phenotype.

To scale up this strong result in a more specific habitat, we applied the same protocol for 502 metabolic models of bacteria found in Tara Ocean Datasets (23). Their pairwise comparison implies considering more than 10^5^ points described by the logarithm of point densities (Fig. 2d). This result confirms the lack of overall relationship between the metabolic niche overlap and the presence-absence of shared reactions for marine prokaryotes. Furthermore, the metabolic niche overlap computation showed several cases of distances near 0 (Fig. 2a-d). This result implies that some metabolic niches were included in others. Due to numerical imprecisions, we stated that an intersection covering more than 999%_0_ of the smallest niche is an inclusion. We used this metric to depict a metabolic niche inclusion graph (Fig. 2e), where nodes are marine bacteria with a node size proportional to the metabolic niche volume, and edges describe the inclusion of a bacteria niche into another. For the sake of clarity, the directed graph follows an isomeric layout driven by the out-degree, and nodes are partitioned into four categories based on the z-score that approximates the capacity to include other metabolic niches. Bacteria known for being associated with a host are depicted in red/bold. Mainly distributed at the bottom of the layout and colored in purple (i.e., metabolic niche included in others), these bacteria show significantly smaller metabolic niche volume (Extended Data Fig. 1). These features indicate less phenotypic plasticity for the bacteria associated with host, potentially following more stable environmental conditions or gene loss (24).

### In-depth study of the metabolic niche flux space of a ubiquitous diatom

To assess the impact of primary metabolism on the metabolic niche and apply our approach beyond prokaryotes, we used one of the most comprehensive and ubiquitous eukaryotic models: the diatom *Phaeodactylum tricornutum* (25). Its metabolic model covers more than 2000 reactions and 1700 metabolites (Supplementary Text 3), which are suited to characterize the chimeric nature of diatom metabolism and incorporate compartmental targeting of biochemical reactions. We investigated the metabolic niche of *P. tricornutum* via a sampling procedure that estimates 10^5^ samples of fluxes distribution that belong to the metabolic niche. This numerical exploration is used to compute pairwise correlations between fluxes. This statistical score emphasizes the relationship between flux variations of two reactions that results from mechanistic dependencies arising from metabolic constraints. Thus, a high absolute correlation value between two reaction fluxes indicates that a change in one of the reaction flux will induce a change in the other reaction flux, whereas not correlated (i.e., R^2^ = 0) fluxes designate mechanistic independence of both reactions in the niche flux space, allowing independent variations of their fluxes. We resumed this exploration in a correlation graph where reactions are vertices and correlation between two reactions is an edge linking the corresponding vertices (Fig. 3a). We assume that the reaction’s importance to sustain the metabolic niche i.e., organism survival, can be shown by the sum of all the correlation involving this particular reaction (as depicted in Fig. 3a by the grey circular bar diagram). We showed that most Calvin-Benson-Bessham (CBB) cycle reactions are essential to the metabolic niche by pointing out reactions belonging to notable metabolic pathways, biologically reassuring for photosynthetic organisms. To be able to get the same kind of results on metabolic pathways, we look at the distribution of its correlation normalized by the number of reactions it is composed of (Fig. 3b). All pathways show a modal distribution around 0, emphasizing that no pathways are entirely dependent on the whole network. Proportionally to others, sulfur and iron show smaller correlations, whereas chlorophyll, carotenoid, amino-acyl-tRNA, glycerolipids, fatty acid biosynthesis, and oxidative stress reactions exhibit higher correlations, as these pathways target energy storage or consumption. Interestingly photorespiration, carbon fixation, TCA, and amino acid metabolism pathways have less high absolute correlation values but still show a normal distribution of the correlations. This result indicates a potential role in acclimating while still having a pivotal role in the organism’s survival.

**Fig. 3.**
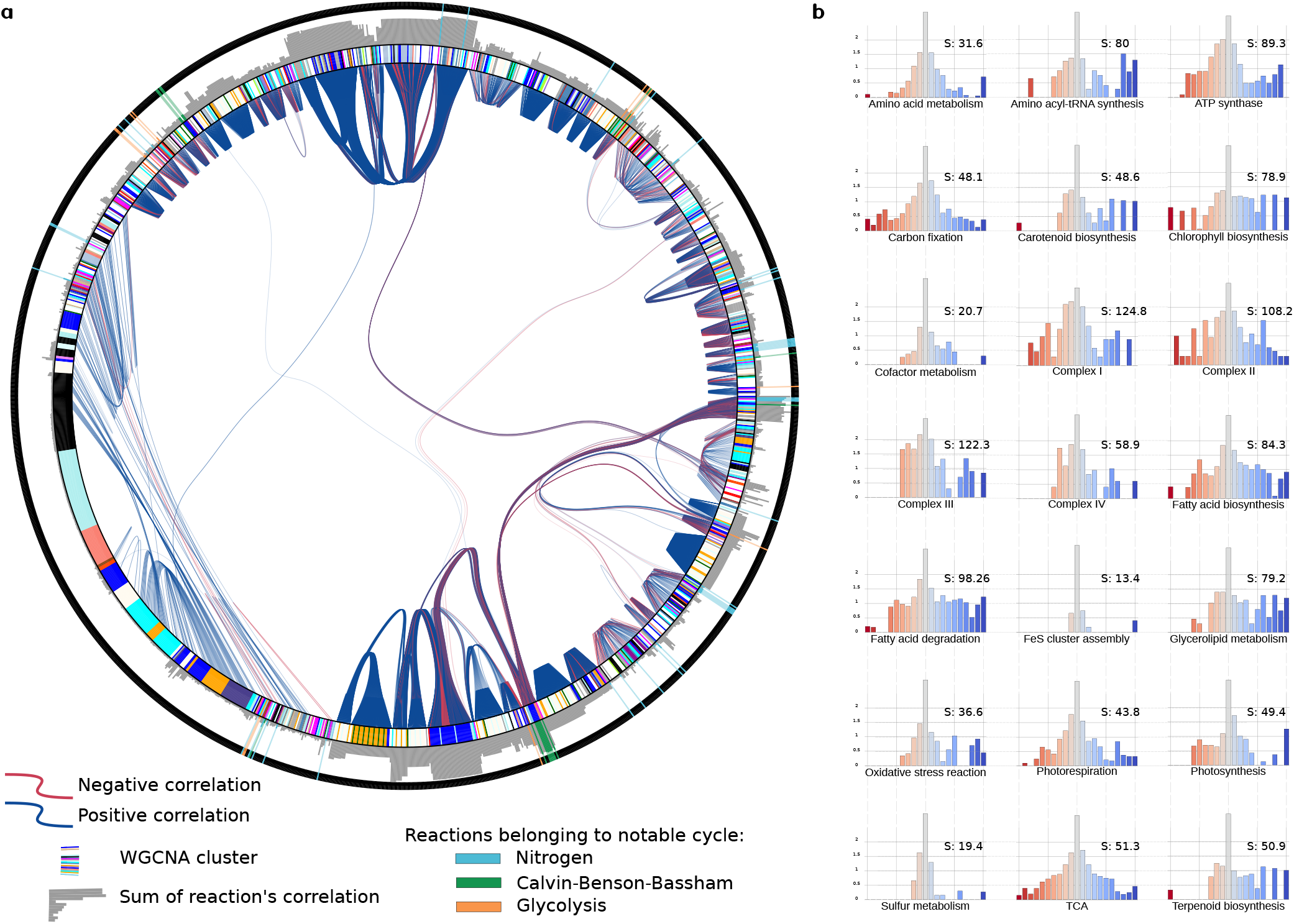
Correlation between Phaeodactylum reactions to support the metabolic niche. **a**, description of most significant dependencies between metabolic reactions. Metabolic reactions are ordered in an outer circle. Reactions associated with nitrogen, Calvin-Benson-Bassham, and glycolysis cycles are emphasized. An edge between reactions corresponds to a correlation (positive or negative) between two fluxes’ reactions above 0.6. The inner-circle describes WGCNA modules to which the gene associated with the reaction belongs. A grey circular bar plot shows the correlation sum of each reaction as a proxy of the reaction essentiality for the metabolic niche. **b** shows essentiality of pathways for the metabolic niche. We represent the correlation histogram normalized by the number of genes present in the pathway for distinct pathways. Y-axes are in log scale. The S score is the sum of the absolute correlation value of the histogram, which estimates pathway essentiality for the metabolic niche.

We can benefit from previous extensive transcriptomic analysis of *P. tricornutum*, in which modules of co-expressed genes over distinct experimental conditions are defined by Weighted Gene Co-expression Network Analysis(26). Because most metabolic reactions are linked to genes that encode for their enzymes, we are able to integrate the information of membership to a module to our analysis (colored ring of Fig. 3a). For some modules, targeted reactions show high pairwise flux correlations, which indicates that the metabolic dependencies could explain the clustering of these genes. Nevertheless, we also see high correlations between modules, indicating metabolic dependencies between different modules (Extended Data Fig. 2a), insights that are not accessible from the standard transcriptomic analysis. Among others, the blue module is of particular interest. It is the most prevalent in the metabolic network. It has the highest intra module correlation sum and shares a high correlation with other modules, emphasizing its importance in the metabolic niche. A previous study (26) showed its implication in several metabolic pathways, mainly the CBB cycle, glycolysis, and fatty acid pathways ( for details, see Fig. S6 of Ait-Mohamed *et al*(26) that depicts the distribution of genes belonging to pathways across all transcriptomic modules). The use of the metabolic niche allows fostering the previous interpretations of these modules. It shows that the module is correlated with reactions directly linked with the TCA pathway (this pathway was essentially associated with Cyan module, explaining in part the connection between these two modules), amino acid metabolism, and chlorophyll biosynthesis (explaining most of the connection with light steel module that encompasses genes associated with chlorophyll and isoprenoid synthesis). Notice that the connection with the dark grey module is primarily due to other pathways not labeled here (Extended Data Fig. 2b).

Finally, projecting the essentiality of genes for the metabolic niche at the chromosome level shows that all chromosomes contribute to the metabolic niche by considering the exact relationship between genes and essential reactions (Fig. 4a). However, extraction of simple statistical parameters from the distribution suggests no particular pattern of gene essentiality for the niche among chromosomes (Fig. 4b). Identifying these islands of essential genes for the metabolic niche vs. segments less constrained paves the way to understand evolutionary processes further.

**Fig. 4.**
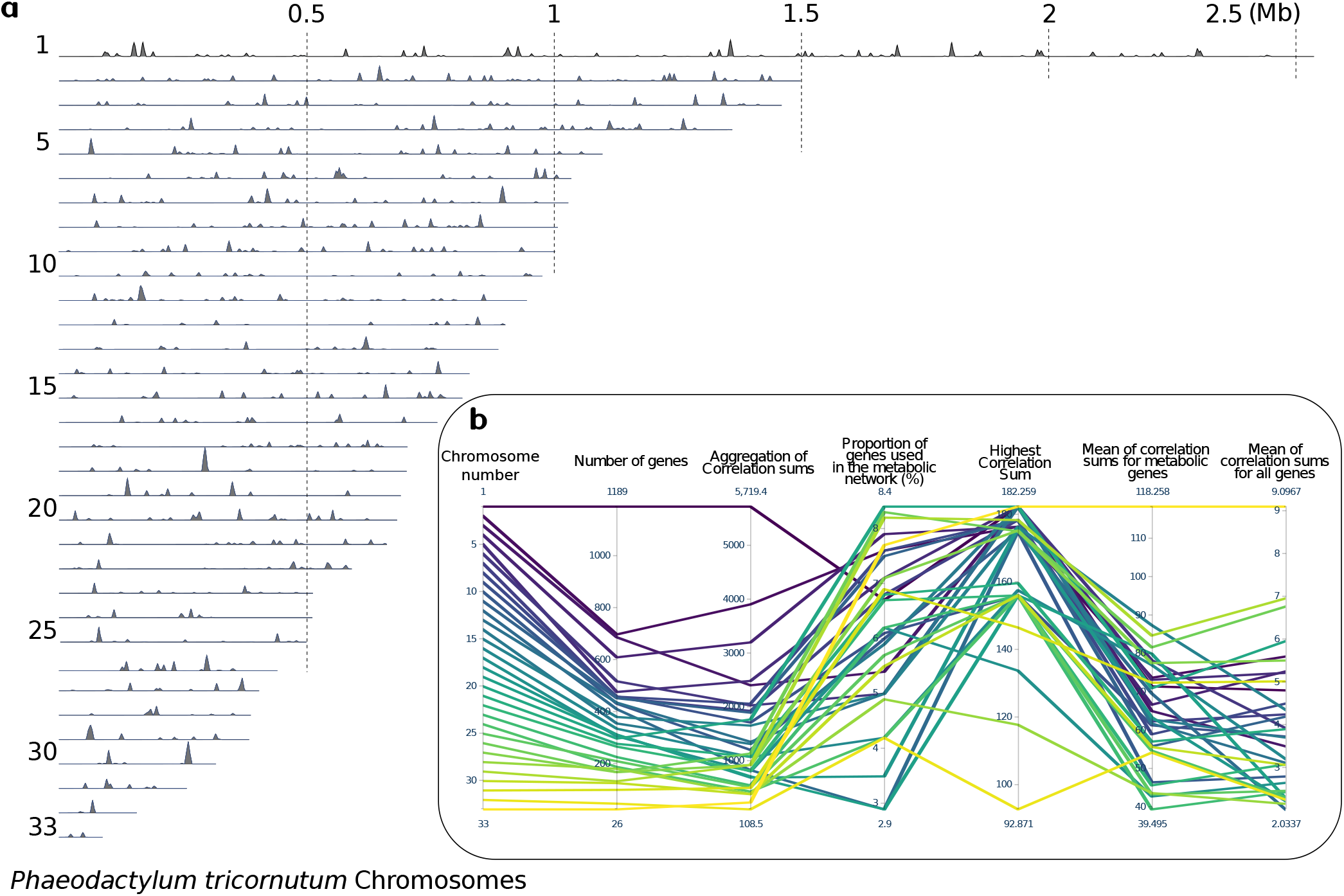
Description of the gene essentiality for the metabolic niche on *Phaedactylum tricornutum*’s genome. **a**, for each 33 diatom’s chromosome, we emphasize genes that encode for a metabolic reaction catalyzer. For each gene, we report the correlation sum of its associated reaction at its genomic position. In addition, **b** describes statistics for each chromosome (i.e., specific color line), such as the total number of genes, the sum of all correlation sums, the proportion of genes from the chromosome involved in the metabolic network, the highest correlation sum found in the chromosome and mean values considering solely genes involved in the metabolic network or all genes.

## Discussion

Our definition of the fundamental niche relies on the recent signs of progress of two recent complementary and productive fields. On the one hand, the genomic composition of organisms and ecosystems is now available via high throughput DNA sequencing fostering the identification of putative functions (10, 27). Notably, a recent study formalizes a metabolic niche space based on the presence-absence of microbe’s metabolic traits, allowing well-defined comparison and clustering of organisms upon metabolic abilities (22). However, it does not investigate the dependency between metabolic capabilities and growth rate (15), putting aside one essential aspect of the niche: the survival of the considered organisms. By doing so, it steps away from Hutchinson’s definition by throwing other niche specificities into the pool (4). On the other hand, systems biology takes advantage of using some of the most efficient computational solvers that enable new descriptions of biological systems phenotypes from networks. The complementarity between these fields allows analyzing genome-scale metabolic models identified from their environment and extrapolating their niche. However, it is worth noting that this models’ identification is a tedious and challenging task (11, 28), and few metabolic models are available compared to the number of annotated genomes. In particular, this study (Fig. 2d,e) considers a fraction of marine prokaryotes, for which metabolic models are available (19), compared to available marine microbial genomes (21). Nevertheless, preliminary metabolic niche exploration is insightful. The metabolic niche inclusion depicts the relative plasticity of bacteria compared to others (Fig. 2e). In particular, it shows that bacteria associated with hosts rely on smaller niches and potentially more stable environmental conditions (Extended Data Fig. 1).

Our metabolic niche approach relies on genome-scale metabolic modeling and is a new formulation of Hutchinson’s fundamental niche. The metabolic niche extracts the main phenotypic characteristics of an organism and considers omics data in the form of a metabolic network as its sole input. Of note, it allows the identification of quantitative traits without the need for parametrizations. For instance, it identified the quantities of oxygen that lead *E. coli* to switch from anaerobic to aerobic growth by just considering its metabolic network (Fig. 1g). This trait description is possible through the sharp abstraction of the internal biochemical machinery, which removes its complexity for emphasizing its effects on growth in the front of nutrient availabilities. From a computational viewpoint, the metabolic niche relies on a formal description of the metabolic solution space. Contrary to other metabolic modeling tools designed for metabolic engineering (used to maximize the growth rate of specific organisms or the production of given metabolites of societal interest), the metabolic niche investigates organismal behaviors considering all kinds of growth conditions, especially sub-optimal. Previous studies advocate for these conditions as more realistic for studying ecological systems or organisms in biotic interactions (29). This statement requires a more exhaustive and computationally challenging exploration of the solution space than extracting small sets of extreme points associated with the maximal growth rate. We proposed the metabolic niche abstraction and the original niche flux space sampling procedure for this purpose.

Formally, the metabolic niche reformulates semi-quantitative knowledge (i.e., presence/absence of genes or relative abundance of gene transcripts) into a quantitative framework to fit the fundamental niche expectation, which is quantitative by nature. This change of abstraction from semi-quantitative to quantitative is a theoretical and computational challenge necessary and recurrent in omics data. The metabolic niche contributes to this general effort by resolving a complicated mathematical problem and assuming the biological system in quasi-steady-state conditions. It is a strong assumption for modeling a biological system in its environment, but it remains coherent with preliminary metabolic engineering studies. Indeed, previous experimental results showed that microbial metabolisms adapt themselves within an hour. Complementary, other constraint-based modeling techniques take benefit from this assumption for simulating an organisms’ adaptation by computing different metabolic fluxes following the evolution of substrates at the minute time-scale (i.e., the systems being at quasi-steady states every minute) (30). These points advocate for the quasi-steady-state assumption and the accuracy of the metabolic niche to investigate the adaptation or acclimation of organisms in environmental conditions.

Interestingly, as shown for *P. tricornutum*, applying a fundamental ecological concept allows integrating a multi-omics data set, which is a fundamental issue in systems biology that often eludes us. In particular, the use of the metabolic niche explains causal dependencies between groups of co-expressed genes. In the case of *P. tricornutum*, it emphasizes the role of the blue module for the diatom survival. Furthermore, when projected on its chromosomes, the metabolic niche concept shows that all chromosomes are necessary for the niche, but not all chromosomic regions are equivalent. This result has implications about the evolution of the organism and how chromosome regions must be more constrained than others for the sake of survival.

Despite biological validations, the metabolic niche remains conceptual and calls for further and extensive bioinformatics applications on large environmental genomic databases such as those provided by the *Tara* Oceans expedition (9). This effort will be necessary to compare our formalism with in situ data about habitat. In addition, the metabolic niche formalization allows precise estimation of the fundamental niche overlap between organisms. However, the work will be colossal as it needs to be performed on more than three thousand metabolic models built from the annotated genomes of reconstructed Metagenome Assembled Genomes (MAGs) (31).

The metabolic niche formalizes the organismal function as a space in which an organism can survive. This new abstraction of the fundamental niche is an addition to other techniques that assess the niche from the presence-absence of omics data (22). In particular, this conceptual study illustrates the need for biological modeling to assess biological phenotype per se as it differs from the sole identification of functional genes. The metabolic niche is thus an essential step towards the design of new omics-trait-based models. It aims to be applied at the organismal and ecosystem levels where we could encompass the whole biological complexity as enclosed in the metagenomic knowledge associated with a superorganism hypothesis (32).

## Methods

### Genome scale metabolic modeling: a general definition

Modeling a metabolic network implies a description of how metabolites are exchanged and transformed within a network. For instance, a general description can take the form of a chemical equation:

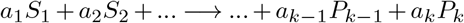

where *S*_*i*_ and *P*_*i*_ depict one metabolite among *k* that are respectively a substrate and a product of the reaction, and *a*_*i*_ its corresponding stoichiometry to satisfy the mass balance law. By definition, substrates or reactants are on the left side of the equation, whereas products are on the right one.

The stoichiometric coefficient *s*_*ij*_ of a metabolite *M*_*i*_ which is involved in a reaction *R*_*j*_ is defined by:

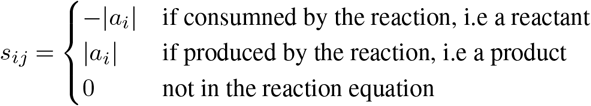

According to kinetic theory, the change over time of the concentration of the metabolite *i* is given by the mass balance equation:

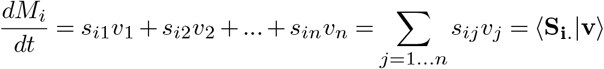

where *v*_*j*_ is the reaction rate or flux associated to reaction *R*_*j*_. Using a vector notation, the above equation can be written as:

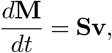

where **M** is a vector that encloses all internal metabolite concentrations, **v** is the flux vector that includes all fluxes *v*_*i*_, and **S** the matrix that stores stoichiometric coefficients and belongs to ℝ^*m,n*^ where *m* is the number of internal metabolites and *n* the number of reactions. Worth noticing, the effect of the temperature on the reaction rate is not taken into account.

This definition of the metabolic network encompasses one particular kind of reactions which are of great interest in the niche definition: the *exchange reactions*, defined as follow:

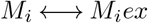

where *M*_*i*_*ex* is a metabolite external to our system. If a metabolite is secreted, the flux through the reaction will be positive (production of *M*_*i*_*ex*). Conversely, a negative flux (production of *M*_*i*_) is an uptake of the metabolite. External metabolites (*M*_*i*_*ex*) are not represented in **M**. In other words, the column of **S** responsible for the exchange reaction will only show a stoichiometric coefficient in the line corresponding to the metabolite in the organism.

Considering that most of the cells are homeostatic, they keep their internal metabolite concentrations as constant as possible. One can thus assume the system at quasi-steady-states (33), leading to:

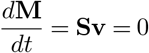

In addition to this system of linear constraints, one also considers the bounds on fluxes that state that no reaction can have an infinite rate. All fluxes must satisfy an inequality like:

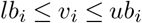

where *ub*_*i*_ represents the upper bound of the flux, meaning the highest rate of the direct reaction, and *lb*_*i*_ represents the lower bound of the flux, i.e. the highest rate of the reverse reaction. One can also encode thermodynamic information by tweaking those bounds. For instance, if the reaction is known to be direct and irreversible, it means that the flux cannot be negative. In that case, the inequality becomes:

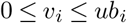

All solutions of v that satisfy these constraints are biochemically accurate. The set of solutions describes a steady-state flux space 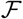 defined by:

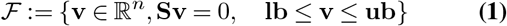

where lb and ub are lower and upper bounds of reaction fluxes. Those values may not be known. In t hat case, a very high value (or a very low value for reversible reactions) is generally used. The corresponding space is then an over-approximation. Each solution of 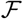 is a point satisfying all chemical and physical constraints in terms of fluxes. Thus, mathematically, points of 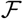 are feasible, as they satisfy all the constraints. Biologically, we will call them functional, as they represent fluxes distribution that allow the organism to survive. We propose herein to investigate the whole set of solutions instead of focusing on one solution proposed by other mechanistic and ultra-parameterized biological modelings.

### General formulation of the metabolic niche

All solutions from 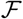 are feasible but represent different phenotypes as the distributions of fluxes indicate different uses of metabolic pathways, or uptakes, or secretions. However, solutions belonging to 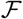 do not form a proper functional niche space. Indeed, two additional constraints must be added to restrain 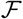 that are described below.

### The survival condition

State-of-the-art metabolic modeling techniques consider a chimeric metabolic reaction, called the biomass reaction (34). The flux through this biomass reaction, *v*_*biomass*_, approximates the growth rate of a given organism and is usually maximized. For instance, in the case of respiration of heterotrophic microbial systems, such a reaction involves more than 25 metabolites. In the genome-scale metabolic framework, growth rate is hence described in mole by mass of dry weight per hour (*mol.gDW*^−1^.*h*^−1^). To survive, an organism must satisfy a flux through this particular reaction above a certain threshold representing its death rate:

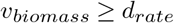

We add this new inequality to the previous set of constraints on flux bounds lb ≤ v ≤ ub. The corresponding solution flux space is noted 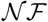 (niche flux space).

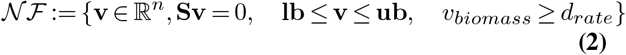

### The environmental space

The metabolic niche space is constrained by the environment, which implies decomposing the flux vector v into two parts: one concerning the exchange reactions (x), and the other concerning the internal reactions (y). Hence, we seek for the set of acceptable x such that there exists a y so that v is in 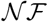. Thus, if among the *n* reactions of the metabolic network, *p* are exchange reactions, we define the metabolic niche by:

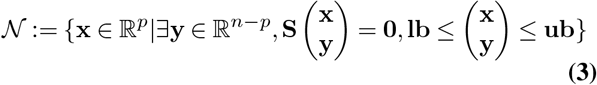

Noting *π*_*ex*_: ℝ*n* ↦ ℝ*p* the orthogonal projection of ℝ*n* onto ℝ*p*, the metabolic niche would be the image of 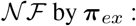 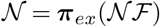.

### Intuition of the projection

One property worth mentioning on 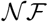 is its topology. This space is convex by definition and also bounded as reactions cannot carry infinite fluxes. Moreover, all its constraints are linear, making it a convex polytope that can be described through the enumeration of its vertices. This representation is called the V-representation (35). Let us call 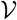 the set of vertices generating 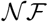. We can write 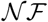 as a convex combination of the vertices in 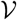, i.e., 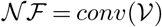. From this description the projection of 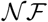 can be done through the projection of each 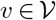and we have:

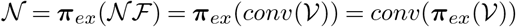

This formulation is well suited for understanding the origin of the niche space in our formalism. However, vertex enumeration is a computationally challenging problem, and its complexity grows exponentially with the number of metabolic reactions involved in the system.

### Metabolic niche computation

This projection of the niche flux space onto the niche space is general as it relies on the seminal definitions of Elton (2) of the fundamental niche that considers all exchange reactions. However, exploring this space is a complicated numerical task. Therefore, we propose to reduce the complexity of the above projection by reformulating the original niche flux space 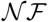, allowing its computation through the resolution of a linear programming problem.

### Polytope formulation

As 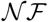 is a polyhedron, one can change its representation from the vertices description to the half-spaces intersection, called the H-representation.

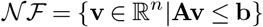

with **A** ∈ ℝ^*q,n*^ and **b** ∈ ℝ^*q*^. **A** is the matrix composed by the vectors defining the half-space constraints (as rows), such that for each row **A**_*i*_., the corresponding constraint is ⟨**A**_*i*_.|**v**⟩ ≤ *b*_*i*_. The new formulation is similar as equation Eq. (2) if **A** and vector **b** are specific matrices defined as follow:

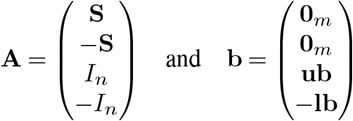

with *I*_*n*_ the identity matrix of ℝ^*n,n*^ and 0_*m*_ the column vector of ℝ^*m*^ with all its components set to 0.

### Projection through multi objective linear programming

Assuming *p* exchange reactions of interest, that represent, axes for the metabolic niche space. Defining the metabolic niche following the formulation of Hutchinson consists in projecting 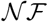 onto the nutrient flux space ℝ^*p*^ defined by those exchange reactions. Formally, it is the polyhedron defined by:

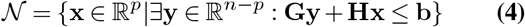

with the two matrices **H** ∈ ℝ^*q,p*^ and **G** ∈ ℝ^*q,n−p*^ being submatrices of **A. H** is composed of the *k* columns corresponding to the exchange reactions; **G** corresponds to the remaining columns taking credit for interior reactions. Biologically speaking, **H** is responsible for the interaction of the organism with its environment, and **G** accounts for the inner mechanism of the organism.

In practice, computing this projection is similar to solve the following Multi-Objective Linear Program (MOLP) (36) for which efficient solvers(37) are available:

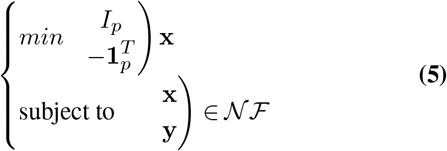

with 1_*p*_ the column vector of ℝ*p* with all its components set to 1. For passing from the MOLP solution to the solution of the projection problem, one only needs to get rid of the last component of the former to get the latter. Computing the niche following the formulation of Hutchinson is therefore equivalent to solve a multi-objective problem in the context of genome-scale metabolic modeling. The detailed pipeline can be found in Appendix A.

### Comparing niches

As p-dimensional volumes, niches can be compared and characterized with different measures. To formally compare such volumes, one can use a pseudo distance based on the Jaccard index (38). The Jaccard index is a similarity measure applied on ensembles, looking at the intersection ratio over the union of the two compared ensembles. The distance is computed as follows:

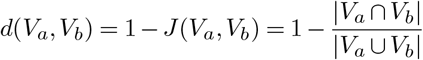

where |.| is an operator measuring the size of the ensemble. For the niche, it is the volume. Biologically speaking, the intersection of two metabolic niches represents all the conditions (fluxes distribution through exchange reactions) that allow both species to survive. The intersection of two p-dimensional volumes is not straightforward, so we developed a method based on metabolic networks. The mathematical and computational framework is explained in Appendix B.

### Exploring niches

The niche flux space investigation emphasizes how the organism allocates its resources and its energy for the sake of its survival. However, the formal investigation of this space is a challenging task that we propose to overcome via the use of OptGP sampler (39) from *COBRApy* (17); see Appendix C for more detail. This technique computes different points that belong to the niche flux space. Each point is a distribution of flux values over all the reactions. Considering fluxes as random variables, we computed pairwise correlations between reactions over the extracted samples to create a weighted correlation graph. It summarizes the metabolic niche’s organization and highlights dependencies between metabolic reactions motivated by its survival. As a final metric, we sum all the correlations associated with one reaction. A high value for a given reaction indicates that this reaction plays a pivotal role as a flux variation of this reaction will imply large changes in several other reactions.

### Genome-scale metabolic networks

We applied the metabolic niche formalism on different metabolic models available in the BiGG database (40). *Escherichia coli str. K-12 substr. MG1655* core is a heterotrophic microbial model organism suitable for bioinformatics benchmarking, comprises 95 reactions of which 20 are exchanges reactions and 72 metabolites. *Phaeodactylum tricornutum CCAP 1055/1* is an ubiquitous eukaryotic organism for which a genome-scale metabolic model is already well described (25). Finally, the metabolic niche was computed on several prokaryotic metabolic models reconstructed with Carveme (19) available at the following repository github.com/cdanielmachado/embl_gems.

## Code availability

The source code for the metabolic niche computation is available at https://gitlab.univ-nantes.fr/aregimbeau/metabolic-niche

## Acknowledgements

We thank the commitment of the following sponsors: CNRS (80 PRIME), and the H2020 project AtlantECO (award number 862923), BiRD bioinformatics facilities, and the Roscoff Bioinformatics platform ABiMS for their computing ressources. Chris Bowler acknowledges the Fonds Français pour l’Environnement Mondial (FFEM), French Government ‘Investissements d’Avenir’ programmes OCEANOMICS (ANR-11-BTBR-0008), FRANCE GENOMIQUE (ANR-10-INBS-09-08), MEMO LIFE (ANR-10-LABX-54), PSL Research University (ANR-11-IDEX-0001-02), the European Research Council (ERC) under the European Union’s Horizon 2020 research and innovation programme (Diatomic; grant agreement No. 835067), and BrownCut (ANR-19-CE20-0020) projects. Marko Budinich and Juan Jose Pierella Karlusich acknowledges postdoctoral funding from FFEM.

## Supplementary Text 1: Computational implementation of the metabolic niche

### Pipeline

From the metabolic network of the considered organism, we identify the exchange reactions (**1**) we want to see as parameters of the niche (whose flux will correspond to the axis of the metabolic niche). With that information, we formulate the problem as a Vector Linear Program (VLP) (**2**), that once solved, results in a list of vertices. The vertices fully characterize the niche as a volume (**3**) in the defined space.

### (1) Identification of Significant exchange reaction

Exchange reactions are explicit from the metabolic network description. However, for the sake of simplicity, the metabolic niche definition requires minimizing the number of exchange reactions. For this purpose, we run a Flux Variability Analysis (FVA) (17) with *COBRApy* that computes both lower and upper bounds of each flux while respecting our constraints. Our constraints are the same as a classical constraint-based model, plus the survival condition, which imposes a minimum flux through the biomass reaction. In our formalism, exchange reactions are considered to have only a reactant and no product. A negative flux describes a consumption by the organism (the metabolite äppearsïn the organism), and a positive flux is a production (the metabolite “disappears”from the organism). Thus every reaction having a negative FVA minimal bound is a reaction responsible for consuming a nutrient. This preliminary check allows us to narrow the number of exchange reactions to consider. For instance, blocked or fixed exchange reactions can bring numerical error in the next step of the niche computation and should be avoided.

### (2) VLP formulation

Once identified, the reactions define a space on which we want to project the niche. From the stoichiometric matrix, the projection matrix, and the reaction bounds, the solver *Bensolve* (37) allows us to solve the problem formulated as follow:

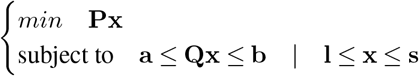

Where **Q** = **S**, a = b = 0 represent the quasi steady states approximation and l and s are respectively lower and upper bound defined in (3). The matrix P is the one defined in (5), with adjustment because the x vector considered in *Bensolve* is our v. The formulation is done through *Benpy* a python wrapping of *Bensolve*. This resolution gives us the upper image of the solution space, which means that we are only interested in the set of vertices returned by the solver and that we need to remove the last component.

### (3) Volume computation

As above, the *Bensolve* solver allows us to get the vertex of the polytope. We then have the V-representation of the polytope. Depending on the network complexity and the size of the projected space, the number of vertices might be vast and difficult to manipulate. To avoid that, one can apply agglomerate clustering to reduce the number of points that simplify the polytope. Once simplified, we can use *lrs* (41) that gives us the volume of the defined polygon.

## Supplementary Text 2: Formalization of the metabolic niche pairwise comparison

The similarity between two niches can be measured through the Jacquard index (38). To do so one need to compute the intersection of a niche pair, as the intersection of multidimensional polytope can be computationally intensive we modify our approach to be able to compute such an intersection.

Let us consider two different species. We then consider the metabolic network and the associated stoichiometric matrices **T** ∈ ℝ^m_T_^,^n_T_^ and **B** ∈ ℝ^m_B_^,^*n*_B_^. To compute the intersection of the two niches, one need to make the assumption that they have exchange reactions in common. Let us note *R*_1_..*R*_*p*_ the *p* exchange reactions we want to consider for the niche computation, and *M*_1_..*M*_*p*_ the *p* corresponding metabolites (for clarity we are omitting here the subscripts *T* and *B* that tell from which organism we are talking about).

We are going to order the matrix **T** and **B**, so that the *p* reactions and metabolites are placed at the top left of the matrix:

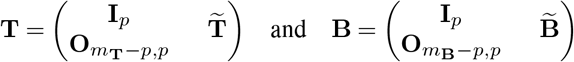

Here the exchange reactions of the two species are distinct axes in the niche space. We need to modify the model so that there is only one set of *p* exchange reactions responsible for the intake of the *p* metabolites of both species. Exchange reaction *i* should look like: *M*_*Bi*_ + *M*_*Ti*_ ↔ *M*_*i*_*ex*. In term of matrix this gives us:

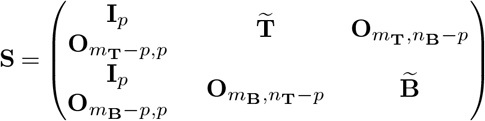

The model has then *m*_*T*_ + *m*_*B*_ metabolites and *n*_*T*_ + *n*_*B*_ –*p* reactions. The first *m*_*T*_ line correspond to the network *T*, and the last *m*_*B*_ lines to the network *B*. The corresponding flux vector x will have its first *p* component responsible for the intake of the *p* metabolites in *T* and *B*, the following *m*_*t*_ – *p* components will be the inner mechanism of *T* that we want to abstract, and the last *m*_*B*_ – *p* lines the one of *B*.

Let us see the implication of such a formalism in term of flux bounds. If we rearranged the bounds order to correspond the matrix T and B defined erlier we have:

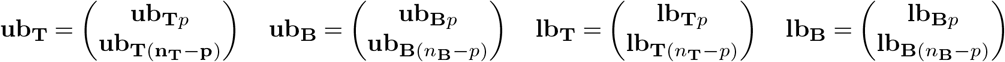

The bounds for the *p* exchange reactions will be the lb_*i*_ = *max*(lb_T*i*_, lb_B*i*_) and ub_*i*_ = *min*(ub_T*i*_, ub_B*i*_). The rest of the bounds are defined by the network that possesses the reaction. Thus we have:

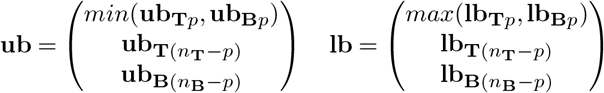

The newly created system can then be formulated as a VLP and the previously described pipeline allows computation of the solution which is the intersection of the two metabolic niches of *T* and *B* computed on the *p* exchange reactions.

## Pairwise comparison of marine prokaryotes

Due to numerical imprecisions or error we made some approximations to make our results more robust. Inclusion where considered when the intersection was covering at least one per mill of the volume of one of the two considered niches. That means, if we consider niche *i* and *j*, with a volume *vol*_*i*_ and *vol*_*j*_, with an intersection *inter* the inclusion was determined if:

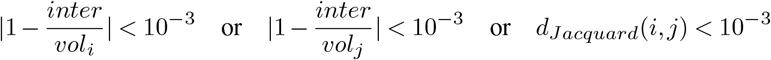

We consider the computation as an error if:

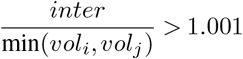

We had 502 species. That means 125751 comparisons. Among those comparisons, 111775 (89%) where computed correctly, 4286 (less than 4% of computed comparisons) results in error because of an intersection that where too big. The rest were either not computed because of an error of the solver, or was taking too long (over than 2 days) during computation (9% of all the comparisons). When computing the inclusion graph we have 47287 edges, an edge is an inclusion. The inclusion relation is transitive, that means that if A include B, and B include C, then A include C. We can applied a transitive closure on the graph.

When we do so we have a graph of 56322 edges. This means that around 10000 comparisons (computed or not) should be inclusion, whereas we got an other results. Half of the newly found edges belongs to not computed comparisons. 15% of them where found in error, which results in 35% of them that are not inclusion in our computations (less than 3% of the computed comparisons). Those errors come from the *Bensolve* solver during computation of the vertices coordinates or from the *Lrs library* during computation of the volume.

## Supplementary Text 3: Sampling of *Phaeodactylum tricornutum* niche space

### Problem description

When formalizing the niche we have a well defined space. The characterization of this space can be done through the interdependences of each pair of reactions. This can be directly measured with the correlation. Method relying on kernel analysis has been proposed to compute correlation between each reaction (42). But this method did not take into account the boundaries of the system, which is for us one key component of our formalism. A way to circumvent this issue is to compute the correlation between reactions through a sampling method. If correctly sampled the niche space allow a quiet accurate computation. But sampling such a high dimensional volume is not an easy thing to do.

### Tools and approximation

We used the OptGP sampler (39) embedded in *COBRApy* to obtain enough points to compute the correlation. We sampled 10^5^ points with a fitting of 10^5^. This allow a convergence of the sampled distribution. We verified the convergence by making 10 batches with the same parameters and look at the variance of each correlation. The sampling has been done with the integration of the survival condition in the model, meaning that the lower bound of the flux through the biomass reaction was set to 0.01. When we sampled the niche space we did not get rid of biomass reactions, whereas a strict sampling of the niche space should be a sampling without the biomass reactions that introduce a bias on the distribution. Indeed the biomass reaction is a constraint, it models the survival of the organism, but the niche definition does not need to have its value, as long as it is above the threshold. Unfortunately the OptGP sampler would requires some heavy modifications to allow this sampling.

### Visualization

Once we have the correlation graph, we need a proper algorithms to visualize it. We used the Graph-Tool library (43). The library implements a hierarchical block structures algorithm(44) which is of great help for module detection in huge graph.

---

**Extended Data Fig. 1.**
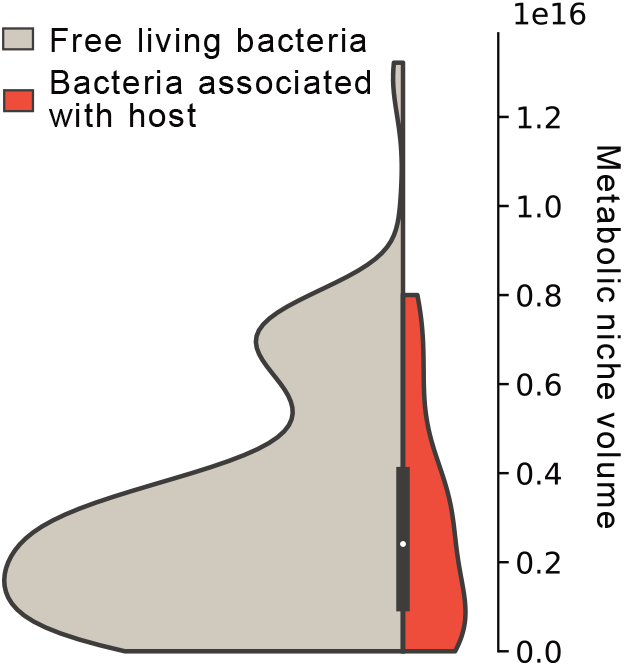
Distribution of metabolic niche volumes for bacteria known for being associated with a host (i.e., red), and bacteria (i.e., in grey) not defined as being associated with host (as stored in PATRIC database)

**Extended Data Fig. 2.**
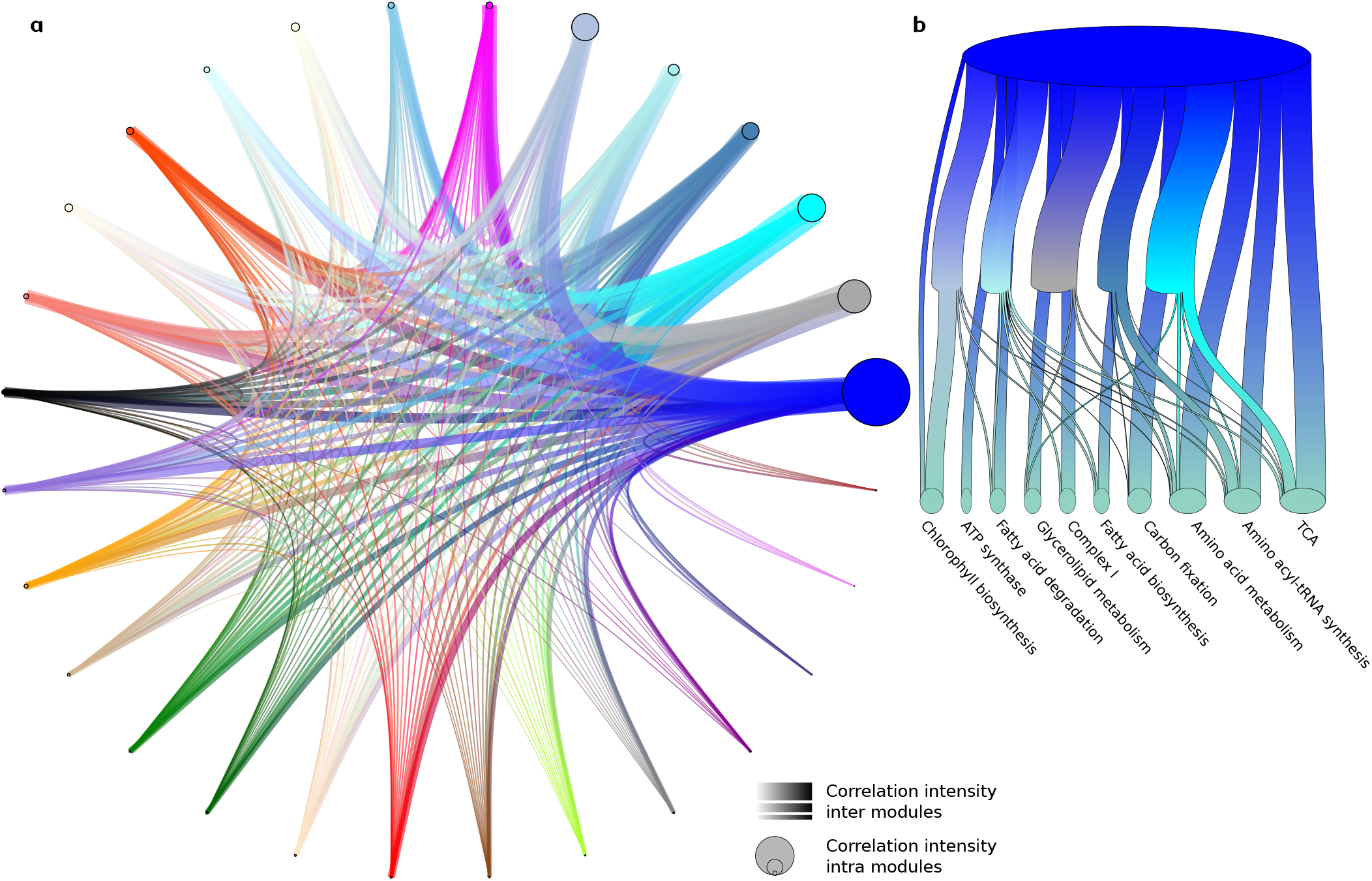
Graph of transcriptomic clusters correlation. On a a general view of the clusters correlations is represented. Each cluster is a vertex, which size depends on the intra-correlation of its genes. The width of the edges represents the value of the correlation sum that is computed between clusters. In b more than 60% of the blue module correlation sum is detailed. A ribbon from the blue ellipse is a correlation that leads to another module and then split into different pathways or directly to a pathway if not related to one of the five represented modules. The intensity of the correlation is proportional to the size of the ribbon.

**Extended Data Fig. 3.**
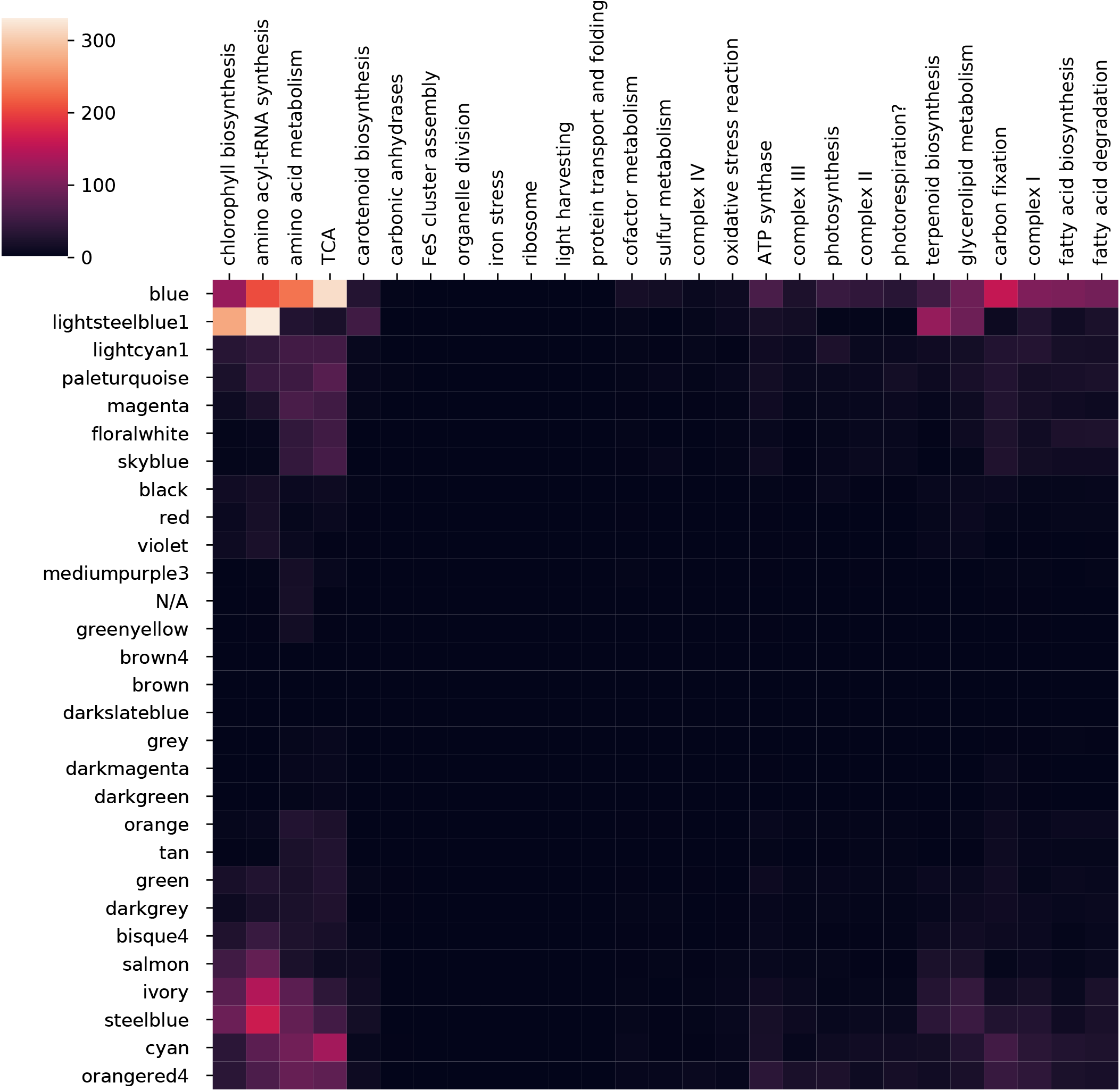
Correlation heatmap of pathways versus modules.

## References

1. Joseph Grinnell. The Niche-Relationships of the California Thrasher. The Auk, 34(4):427–433, 1917. ISSN 0004-8038. doi: 10.2307/4072271.

2. Charles S. (Charles Sutherland) Elton. Animal ecology. New York, Macmillan Co., 1927.

3. G. E. Hutchinson. Concluding Remarks. Cold Spring Harbor Symposia on Quantitative Biology, 22(0):415–427, January 1957. ISSN 0091-7451, 1943-4456. doi: 10.1101/SQB.1957.022.01.039.

4. Greg J. McInerny and Rampal S. Etienne. Ditch the niche – is the niche a useful concept in ecology or species distribution modelling? Journal of Biogeography, 39(12):2096–2102, 2012. ISSN 1365-2699. doi: 10.1111/jbi.12033.

5. Jessica L Green, Brendan J M Bohannan, and Rachel J Whitaker. Microbial Biogeography: From Taxonomy to Traits. Science, 320(5879):1039–1043, may 2008. ISSN 0036-8075. doi: 10.1126/science.1153475.

6. Hiroaki Kitano. Systems Biology: A Brief Overview. Science, 295(5560):1662–1664, mar 2002. ISSN 00368075. doi: 10.1126/science.1069492.

7. Andrew R. Joyce and Bernhard Palsson. The model organism as a system: Integrating‘omics’ data sets. Nature Reviews Molecular Cell Biology, 7(3):198–210, mar 2006. ISSN 14710072. doi: 10.1038/nrm1857.

8. Jennifer Levering, Christopher L. Dupont, Andrew E. Allen, Bernhard O. Palsson, and Karsten Zengler. Integrated Regulatory and Metabolic Networks of the Marine Diatom Phaeo-dactylum tricornutum Predict the Response to Rising CO 2 Levels. mSystems, 2(1), 2017. ISSN 2379-5077. doi: 10.1128/msystems.00142-16.

9. Shinichi Sunagawa, Silvia G. Acinas, Peer Bork, Chris Bowler, Silvia G. Acinas, Marcel Babin, Peer Bork, Emmanuel Boss, Chris Bowler, Guy Cochrane, Colomban de Vargas, Michael Follows, Gabriel Gorsky, Nigel Grimsley, Lionel Guidi, Pascal Hingamp, Daniele Iudicone, Olivier Jaillon, Stefanie Kandels, Lee Karp-Boss, Eric Karsenti, Magali Lescot, Fabrice Not, Hiroyuki Ogata, Stéphane Pesant, Nicole Poulton, Jeroen Raes, Christian Sardet, Mike Sieracki, Sabrina Speich, Lars Stemmann, Matthew B. Sullivan, Shinichi Sunagawa, Patrick Wincker, Damien Eveillard, Gabriel Gorsky, Lionel Guidi, Daniele Iudicone, Eric Karsenti, Fabien Lombard, Hiroyuki Ogata, Stephane Pesant, Matthew B. Sullivan, Patrick Wincker, and Colomban de Vargas. Tara Oceans: towards global ocean ecosystems biology. Nature Reviews Microbiology, 18(8):428–445, 2020. ISSN 17401534. doi: 10.1038/s41579-020-0364-5.

10. V. J. Coles, M. R. Stukel, M. T. Brooks, A. Burd, B. C. Crump, M. A. Moran, J. H. Paul, B. M. Satinsky, P. L. Yager, B. L. Zielinski, and R. R. Hood. Ocean biogeochemistry modeled with emergent trait-based genomics. Science, 358(6367):1149–1154, dec 2017. ISSN 10959203. doi: 10.1126/science.aan5712.

11. Ines Thiele and Bernhard Ø Palsson. A protocol for generating a high-quality genome-scale metabolic reconstruction. Nature protocols, 5(1):93–121, 2010. doi: 10.1038/nprot.2009.203.

12. Jeffrey D. Orth, Ines Thiele, and Bernhard O. Palsson. What is flux balance analysis? Nature Biotechnology, 28(3):245–248, mar 2010. ISSN 10870156. doi: 10.1038/nbt.1614.

13. Aleksej Zelezniak, Sergej Andrejev, Olga Ponomarova, Daniel R. Mende, Peer Bork, and Kiran Raosaheb Patil. Metabolic dependencies drive species co-occurrence in diverse microbial communities. Proceedings of the National Academy of Sciences of the United States of America, 112(20):6449–6454, may 2015. ISSN 10916490. doi: 10.1073/pnas.1421834112.

14. Csaba Pál and Balázs Papp. Evolution of complex adaptations in molecular systems. Nature Ecology Evolution, 1(8):1084–1092, aug 2017. ISSN 2397-334X. doi: 10.1038/s41559-017-0228-1.

15. Sarah R. Smith, Chris L. Dupont, James K. McCarthy, Jared T. Broddrick, Miroslav Oborník, Aleš Horák, Zoltán Füssy, Jaromír Cihlářr, Sabrina Kleessen, Hong Zheng, John P. McCrow, Kim K. Hixson, Wagner L. Araújo, Adriano Nunes-Nesi, Alisdair Fernie, Zoran Nikoloski, Bernhard O. Palsson, and Andrew E. Allen. Evolution and regulation of nitrogen flux through compartmentalized metabolic networks in a marine diatom. Nature Communications, 10(1): 4514–4552, 2019. ISSN 20411723. doi: 10.1038/s41467-019-12407-y.

16. Antoine Guisan and Wilfried Thuiller. Predicting species distribution: Offering more than simple habitat models. Ecology Letters, 8(9):993–1009, sep 2005. ISSN 1461023X. doi: 10.1111/j.1461-0248.2005.00792.x.

17. Laurent Heirendt, Sylvain Arreckx, Thomas Pfau, Sebastián N. Mendoza, Anne Richelle, Almut Heinken, Hulda S. Haraldsdóttir, Jacek Wachowiak, Sarah M. Keating, Vanja Vlasov, Stefania Magnusdóttir, Chiam Yu Ng, German Preciat, Alise Žagare, Siu H.J. Chan, Maike K. Aurich, Catherine M. Clancy, Jennifer Modamio, John T. Sauls, Alberto Noronha, Aarash Bordbar, Benjamin Cousins, Diana C. El Assal, Luis V. Valcarcel, Iñigo Apaolaza, Susan Ghaderi, Masoud Ahookhosh, Marouen Ben Guebila, Andrejs Kostromins, Nicolas Sompairac, Hoai M. Le, Ding Ma, Yuekai Sun, Lin Wang, James T. Yurkovich, Miguel A.P. Oliveira, Phan T. Vuong, Lemmer P. El Assal, Inna Kuperstein, Andrei Zinovyev, H. Scott Hinton, William A. Bryant, Francisco J. Aragón Artacho, Francisco J. Planes, Egils Stalidzans, Alejandro Maass, Santosh Vempala, Michael Hucka, Michael A. Saunders, Costas D. Maranas, Nathan E. Lewis, Thomas Sauter, Bernhard Palsson, Ines Thiele, and Ronan M.T. Fleming. Creation and analysis of biochemical constraint-based models using the COBRA Toolbox v.3.0. Nature Protocols, 14(3):639–702, 2019. ISSN 17502799. doi: 10.1038/s41596-018-0098-2.

18. Sandra Díaz, Jens Kattge, Johannes H.C. Cornelissen, Ian J. Wright, Sandra Lavorel, Stéphane Dray, Björn Reu, Michael Kleyer, Christian Wirth, I. Colin Prentice, Eric Garnier, Gerhard Bönisch, Mark Westoby, Hendrik Poorter, Peter B. Reich, Angela T. Moles, John Dickie, Andrew N. Gillison, Amy E. Zanne, Jérôme Chave, S. Joseph Wright, Serge N. Sheremet Ev, Hervé Jactel, Christopher Baraloto, Bruno Cerabolini, Simon Pierce, Bill Shipley, Donald Kirkup, Fernando Casanoves, Julia S. Joswig, Angela Günther, Valeria Falczuk, Nadja Rüger, Miguel D. Mahecha, and Lucas D. Gorné. The global spectrum of plant form and function. Nature, 529(7585):167–171, 2016. ISSN 14764687. doi: 10.1038/nature16489.

19. Daniel Machado, Sergej Andrejev, Melanie Tramontano, and Kiran Raosaheb Patil. Fast automated reconstruction of genome-scale metabolic models for microbial species and communities. Nucleic Acids Research, 46(15):7542–7553, 06 2018. ISSN 0305-1048. doi: 10.1093/nar/gky537.

20. James J. Davis, Alice R. Wattam, Ramy K. Aziz, Thomas Brettin, Ralph Butler, Rory M. Butler, Philippe Chlenski, Neal Conrad, Allan Dickerman, Emily M. Dietrich, Joseph L. Gabbard, Svetlana Gerdes, Andrew Guard, Ronald W. Kenyon, Dustin MacHi, Chunhong Mao, Dan Murphy-Olson, Marcus Nguyen, Eric K. Nordberg, Gary J. Olsen, Robert D. Olson, Jamie C. Overbeek, Ross Overbeek, Bruce Parrello, Gordon D. Pusch, Maulik Shukla, Chris Thomas, Margo Vanoeffelen, Veronika Vonstein, Andrew S. Warren, Fangfang Xia, Dawen Xie, Hyunseung Yoo, and Rick Stevens. The PATRIC Bioinformatics Resource Center: Expanding data and analysis capabilities. Nucleic Acids Research, 48(D1):D606–D612, oct 2020. ISSN 13624962. doi: 10.1093/nar/gkz943.

21. Stilianos Louca, Laura Wegener Parfrey, and Michael Doebeli. Decoupling function and taxonomy in the global ocean microbiome. Science, 353(6305):1272–1277, September 2016. ISSN 0036-8075. doi: 10.1126/science.aaf4507.

22. Ashkaan K. Fahimipour, and Thilo Gross. Mapping the bacterial metabolic niche space. Nature Communications, 11(1):415–418, 2020. ISSN 20411723. doi: 10.1038/s41467-020-18695-z.

23. Caroline Vernette, Nicolas Henry, Julien Lecubin, Colomban Vargas, Pascal Hingamp, and Magali Lescot. The Ocean barcode atlas: A web service to explore the biodiversity and biogeography of marine organisms. Molecular Ecology Resources, 21(4):1347–1358, may 2021. ISSN 1755-098X. doi: 10.1111/1755-0998.13322.

24. Alix Mas, Shahrad Jamshidi, Yvan Lagadeuc, Damien Eveillard, and Philippe Vandenkoornhuyse. Beyond the Black Queen Hypothesis. The ISME Journal, 10(9):2085–2091, sep 2016. ISSN 1751-7362. doi: 10.1038/ismej.2016.22.

25. Jared T. Broddrick, Niu Du, Sarah R. Smith, Yoshinori Tsuji, Denis Jallet, Maxwell A. Ware, Graham Peers, Yusuke Matsuda, Chris L. Dupont, B. Greg Mitchell, Bernhard O. Palsson, and Andrew E. Allen. Cross-compartment metabolic coupling enables flexible photoprotective mechanisms in the diatom Phaeodactylum tricornutum. New Phytologist, 222(3): 1364–1379, may 2019. ISSN 14698137. doi: 10.1111/nph.15685.

26. Ouardia Ait-Mohamed, Anna M. G. Novák Vanclová, Nathalie Joli, Yue Liang, Xue Zhao, Auguste Genovesio, Leila Tirichine, Chris Bowler, and Richard G. Dorrell. Phaeonet: A holistic rnaseq-based portrait of transcriptional coordination in the model diatom phaeodactylum tricornutum. Frontiers in Plant Science, 11:1522, 2020. ISSN 1664-462X. doi: 10.3389/fpls.2020.590949.

27. Jeroen Raes and Peer Bork. Molecular eco-systems biology: Towards an understanding of community function. Nature Reviews Microbiology, 6(9):693–699, sep 2008. ISSN 17401526. doi: 10.1038/nrmicro1935.

28. Christian Lieven, Moritz E. Beber, Brett G. Olivier, Frank T. Bergmann, Meric Ataman, Parizad Babaei, Jennifer A. Bartell, Lars M. Blank, Siddharth Chauhan, Kevin Correia, Christian Diener, Andreas Dräger, Birgitta E. Ebert, Janaka N. Edirisinghe, José P. Faria, Adam M. Feist, Georgios Fengos, Ronan M. T. Fleming, Beatriz García-Jiménez, Vassily Hatzimanikatis, Wout van Helvoirt, Christopher S. Henry, Henning Hermjakob, Markus J. Herrgård, Ali Kaafarani, Hyun Uk Kim, Zachary King, Steffen Klamt, Edda Klipp, Jasper J. Koehorst, Matthias König, Meiyappan Lakshmanan, Dong-Yup Lee, Sang Yup Lee, Sunjae Lee, Nathan E. Lewis, Filipe Liu, Hongwu Ma, Daniel Machado, Radhakrishnan Mahadevan, Paulo Maia, Adil Mardinoglu, Gregory L. Medlock, Jonathan M. Monk, Jens Nielsen, Lars Keld Nielsen, Juan Nogales, Intawat Nookaew, Bernhard O. Palsson, Jason A. Papin, Kiran R. Patil, Mark Poolman, Nathan D. Price, Osbaldo Resendis-Antonio, Anne Richelle, Isabel Rocha, Benjamín J. Sánchez, Peter J. Schaap, Rahuman S. Malik Sheriff, Saeed Shoaie, Nikolaus Sonnenschein, Bas Teusink, Paulo Vilaça, Jon Olav Vik, Judith A. H. Wodke, Joana C. Xavier, Qianqian Yuan, Maksim Zakhartsev, and Cheng Zhang. MEMOTE for standardized genome-scale metabolic model testing. Nature Biotechnology, 38(3):272–276, March 2020. ISSN 1087-0156. doi: 10.1038/s41587-020-0446-y.

29. Marko Budinich, Jérémie Bourdon, Abdelhalim Larhlimi, and Damien Eveillard. A multi-objective constraint-based approach for modeling genome-scale microbial ecosystems. PLoS ONE, 12(2):e0171744, 2017. ISSN 19326203. doi: 10.1371/journal.pone.0171744.

30. Ana Bulovicć, Stephan Fischer, Marc Dinh, Felipe Golib, Wolfram Liebermeister, Christian Poirier, Laurent Tournier, Edda Klipp, Vincent Fromion, and Anne Goelzer. Automated generation of bacterial resource allocation models. Metabolic Engineering, 55(June):12–22, 2019. ISSN 10967184. doi: 10.1016/j.ymben.2019.06.001.

31. Guillem Salazar, Lucas Paoli, Adriana Alberti, Jaime Huerta-Cepas, Hans Joachim Ruscheweyh, Miguelangel Cuenca, Christopher M. Field, Luis Pedro Coelho, Corinne Cruaud, Stefan Engelen, Ann C. Gregory, Karine Labadie, Claudie Marec, Eric Pelletier, Marta Royo-Llonch, Simon Roux, Pablo Sánchez, Hideya Uehara, Ahmed A. Zayed, Georg Zeller, Margaux Carmichael, Céline Dimier, Joannie Ferland, Stefanie Kandels, Marc Picheral, Sergey Pisarev, Julie Poulain, Silvia G. Acinas, Marcel Babin, Peer Bork, Emmanuel Boss, Chris Bowler, Guy Cochrane, Colomban de Vargas, Michael Follows, Gabriel Gorsky, Nigel Grimsley, Lionel Guidi, Pascal Hingamp, Daniele Iudicone, Olivier Jaillon, Stefanie Kandels-Lewis, Lee Karp-Boss, Eric Karsenti, Fabrice Not, Hiroyuki Ogata, Stephane Pesant, Nicole Poulton, Jeroen Raes, Christian Sardet, Sabrina Speich, Lars Stemmann, Matthew B. Sullivan, Shinichi Sunagawa, and Patrick Wincker. Gene Expression Changes and Community Turnover Differentially Shape the Global Ocean Metatranscriptome. Cell, 179(5): 1068–1083.e21, 2019. ISSN 10974172. doi: 10.1016/j.cell.2019.10.014.

32. Kevin Liautaud, Egbert H van Nes, Matthieu Barbier, Marten Scheffer, and Michel Loreau. Superorganisms or loose collections of species? A unifying theory of community patterns along environmental gradients. Ecology Letters, 22(8):1243–1252, 2019. doi: 10.1111/ele.13289.

33. Amit Varma and Bernhard O. Palsson. Metabolic flux balancing: Basic concepts, scientific and practical use. Nat Biotechnol, 12(10):994–998, oct 1994. ISSN 0733222X. doi: 10.1038/nbt1094-994.

34. Adam M. Feist, and Bernhard O. Palsson. The biomass objective function. Current Opinion in Microbiology, 13(3):344–349, jun 2010. ISSN 13695274. doi: 10.1016/j.mib.2010.03.003.

35. Günter M. Ziegler. Faces of Polytopes. In Günter M. Ziegler, editor, Lectures on Polytopes: Updated Seventh Printing of the First Edition, Graduate Texts in Mathematics, pages 51–76. Springer, New York, NY, 1995. ISBN 978-1-4613-8431-1. doi: 10.1007/978-1-4613-8431-1_2.

36. Andreas Löhne and Benjamin Weißing. Equivalence between polyhedral projection, multiple objective linear programming and vector linear programming. Mathematical Methods of Operations Research, 84(2):411–426, October 2016. ISSN 1432-5217. doi: 10.1007/s00186-016-0554-0.

37. Andreas Löhne and Benjamin Weißing. The vector linear program solver Bensolve – notes on theoretical background. European Journal of Operational Research, 260(3):807–813, 2017. ISSN 03772217. doi: 10.1016/j.ejor.2016.02.039.

38. A. Conci and C. S. Kubrusly. Distance Between Sets - A survey. Advances in Mathematical Sciences and Applications, 2017.

39. Wout Megchelenbrink, Martijn Huynen, and Elena Marchiori. optGpSampler: An improved tool for uniformly sampling the solution-space of genome-scale metabolic networks. PLoS ONE, 9(2), 2014. ISSN 19326203. doi: 10.1371/journal.pone.0086587.

40. Zachary A. King, Justin Lu, Andreas Dräger, Philip Miller, Stephen Federowicz, Joshua A. Lerman, Ali Ebrahim, Bernhard O. Palsson, and Nathan E. Lewis. BiGG Models: A platform for integrating, standardizing and sharing genome-scale models. Nucleic Acids Research, 44(D1):D515–D522, jan 2016. ISSN 13624962. doi: 10.1093/nar/gkv1049.

41. David Avis. A Revised Implementation of the Reverse Search Vertex Enumeration Algorithm. In Polytopes — Combinatorics and Computation, volume c, pages 177–198. Birkhäuser Basel, Basel, 2000. doi: 10.1007/978-3-0348-8438-9_9.

42. Jean Marc Schwartz, Hiroaki Otokuni, Tatsuya Akutsu, and Jose C. Nacher. Probabilistic controllability approach to metabolic fluxes in normal and cancer tissues. Nature Communications, 10(1):1–9, 2019. ISSN 20411723. doi: 10.1038/s41467-019-10616-z.

43. Tiago P. Peixoto. The graph-tool python library. figshare, 2014. doi: 10.6084/m9.figshare.1164194.

44. Tiago P. Peixoto. Hierarchical block structures and high-resolution model selection in large networks. Phys. Rev. X, 4(1):011047, 2014. doi: 10.1103/PhysRevX.4.011047. Publisher: American Physical Society.

